# Controlled decorin delivery from injectable microgels promotes scarless vocal fold repair

**DOI:** 10.1101/2025.06.25.661429

**Authors:** Ryan M. Friedman, Elizabeth A. Brown, Hannah M. Bonelli, Yashna Gupta, Matthew R. Aronson, Kendra McDaid, Hannah M. Oh, Eiman Abu Bandora, Karen B. Zur, Riccardo Gottardi

## Abstract

Vocal fold (VF) scarring is a leading cause of poor voice, yet no therapies exist to prevent its progression. Current treatments, such as intracordal steroid injections, offer limited efficacy and carry significant off-target toxicities. To identify targeted anti-scarring strategies, we performed transcriptomics of human VF myofibroblasts, the cellular drivers of VF scarring, and identified the proteoglycan decorin (DCN) as downregulated in activated myofibroblasts. We also show a time-dependent decrease in DCN during fibrotic wound healing in a preclinical rat model of VF scarring. Administration of DCN suppressed VF myofibroblast activation by reducing pro-fibrotic gene expression, α-smooth muscle actin (α-SMA) levels, and cell contractility. DCN was encapsulated in hyaluronic acid microgels for sustained protein release for 3–4 weeks. In a rat model of VF scarring, DCN-loaded microgels prevented hallmark features of scarring, including collagen deposition and myofibroblast activation. These findings highlight DCN as a promising therapeutic and provide a sustained delivery platform with translational potential against VF scarring.

**Teaser:** Controlled decorin drug delivery from hyaluronic acid microgels attenuates myofibroblast activation to prevent vocal fold scarring, offering a translational strategy to improve post-injury voice outcomes.

## Introduction

Voicing is a complex physiologic process important for daily communication, knowledge transfer, cultural expression, and social connectivity. Phonation requires proper vibration of the vocal folds (VFs), and damage to these tissues leads to poor vocal quality. Nearly 30% of the general population is affected by voice disorders during their lifetime (*1*), resulting in an annual economic burden of $13 billion (*2*). Patients experience emotional distress (*3*), decreased social engagement (*4*), and an overall reduction in quality of life (*5–7*). The greatest refractory cause of dysphonia, broadly defined as poor voice, after laryngeal injury is vocal fold scarring (*8, 9*). Scar formation typically follows severe upper airway trauma or iatrogenic injuries to the VF epithelium and underlying lamina propria (*8*). After injury, immune cells infiltrate the wound bed and secrete pro-inflammatory cytokines, such as transforming growth factor-β (TGF-β) isoforms, to promote tissue repair (*10, 11*). In response to these secreted signals, resident VF fibroblasts undergo a fibroblast-to-myofibroblast transition (*12*). While myofibroblasts are required for wound closure and VF repair, persistent myofibroblast activation results in the over-accumulation of a collagen-rich extracellular matrix (ECM). This process leads to tissue stiffening and scar formation (*8*). Altogether, these changes disrupt normal mucosal wave propagation, and as a result, patients experience vocal hoarseness, vocal fatigue, and poor voice control (*13*).

Intracordal steroid injections administered directly into the scar tissue provide broad anti-inflammatory effects and are the most common strategy for early management of VF scars (*9, 14, 15*). However, steroids do not target any specific pro-fibrotic mechanisms implicated in the pathology of VF scarring and are only intended to alleviate symptoms of dysphonia (*16*). Since steroids fail to resolve ongoing fibrosis, any marginal improvement to acoustic parameters is temporary, with success rates dropping to as low as 57% after 6 months. (*17*). Over time, VF scarring leads to glottic insufficiency, i.e., the incomplete closure of the VFs during phonation, due to prolonged tissue contraction and scar-associated tissue atrophy. Surgical approaches, such as injection medialization laryngoplasty (IML), are often performed as an alternative to intracordal steroid injections. In IML a biomaterial is directly injected into the paraglottic space, lateral to the VF lamina propria or underlying vocalis muscle (*18*), to medialize the VFs and compensate glottic insufficiency. However, IML is not disease-modifying and does not promote scarless tissue repair. Currently, there are no clinically-available antifibrotic agents that halt ongoing scar formation or restore healthy tissue structure (*13*).

The development and success of new VF scarring therapeutics is further complicated by significant drug delivery challenges in the upper airway. Intracordal VF injections offer the most straightforward route of administration and bypass the VF mucus layer, which limits drug absorption. However, this approach is associated with an increased risk of surgical complications due to the complex anatomy of the upper airway and the relatively small size of VFs (*19*). In fact, repeated injections are often necessary because of the short half-life of steroids, which leads to safety concerns and increases the risk of off-target toxicities such as VF atrophy, mucosal gland atrophy, and epithelial thinning (*14, 16, 20*). To overcome these limitations, controlled drug delivery systems may be engineered to improve the efficacy and safety profile of new antifibrotic molecules.

This study aimed to determine genes that regulate VF myofibroblast activation, uncovering potential therapeutic targets for anti-scarring strategies, and to engineer a biomaterial platform for controlled drug delivery to the VF. To identify therapeutic candidates, we compared the transcriptome of activated human VF myofibroblasts to resting VF fibroblast controls. Decorin (DCN), a small leucine-rich proteoglycan implicated in VF development and VF ECM organization (*21*), was one of the most significantly downregulated genes in myofibroblasts *in vitro* and was depleted in scarred vocal rat VFs *in vivo*. We then established DCN as a strong antifibrotic protein, able to reduce VF myofibroblast activation. Based on these observations, we hypothesized that the addition of DCN is sufficient to attenuate scar formation after direct VF injury by reducing myofibroblast activation. We engineered an injectable hyaluronic acid microgel platform capable of sustained DCN delivery over 3 4 weeks. In a rat model of VF scarring, intracordal injection of DCN-loaded microgels suppressed fibrotic wound healing and supported scarless tissue repair, supporting the potential of this platform as a mechanism-based, targeted anti-fibrotic therapy.

## Results

### Decorin is downregulated in vocal fold myofibroblasts and in scarred vocal folds

Human VF fibroblasts, the primary cell type in the VF lamina propria (*22*), regulate fibrogenesis, scar formation, and tissue remodeling after injury (*23–25*). To better understand the progression of VF scarring, we performed transcriptomics on human VF fibroblasts treated with TGF-β1 to induce myofibroblast activation, which were compared to unstimulated, resting VF fibroblast controls. Bulk RNA sequencing revealed higher similarity within experimental groups than within donor groups, suggesting that TGF-β1 strongly affects the gene expression of VF fibroblasts *in vitro* (Fig. 1A). Unsupervised clustering of all samples by their differentially expressed genes (DEGs) shows that the transcriptome of VF myofibroblasts is uniquely distinct from VF fibroblasts (Fig. 1B). Gene set enrichment analysis (GSEA) using Gene Ontology (GO) annotations was used to connect transcriptional profiles to functional biological outputs (*26, 27*). As expected, GSEA highlights positive enrichment for many pathways involving either myofibroblast activation or ECM secretion and organization (Fig. 1C), confirming that TGF-β1 successfully induced VF myofibroblast activation *in vitro*. Since many pathways involving the ECM were positively enriched, we also carried out GSEA with Matrisome annotations. This approach revealed 128 downregulated and 148 upregulated DEGs in the core matrisome or matrisome-associated sets (Fig. 1D). Among core matrisome genes, *DCN*, encoding for the DCN protein, is the most significantly downregulated gene (Table S1). *DCN* is also the third-most significantly downregulated gene in the entire transcriptome of VF myofibroblasts (Fig. 1E). Notably, genes encoding for other small-leucine rich proteoglycans in the same family and class as DCN (*28*) are not downregulated in our dataset (Fig. S1A). RT-qPCR results confirm our transcriptomics data and suggest that *DCN* is downregulated in VF myofibroblasts with increased gene expression of the myofibroblast marker *ACTA2* (Fig. 1F). In addition, both intracellular and secreted DCN protein abundance is significantly reduced in VF myofibroblasts (Fig. S1B, Fig. 1G).

**Fig. 1.**
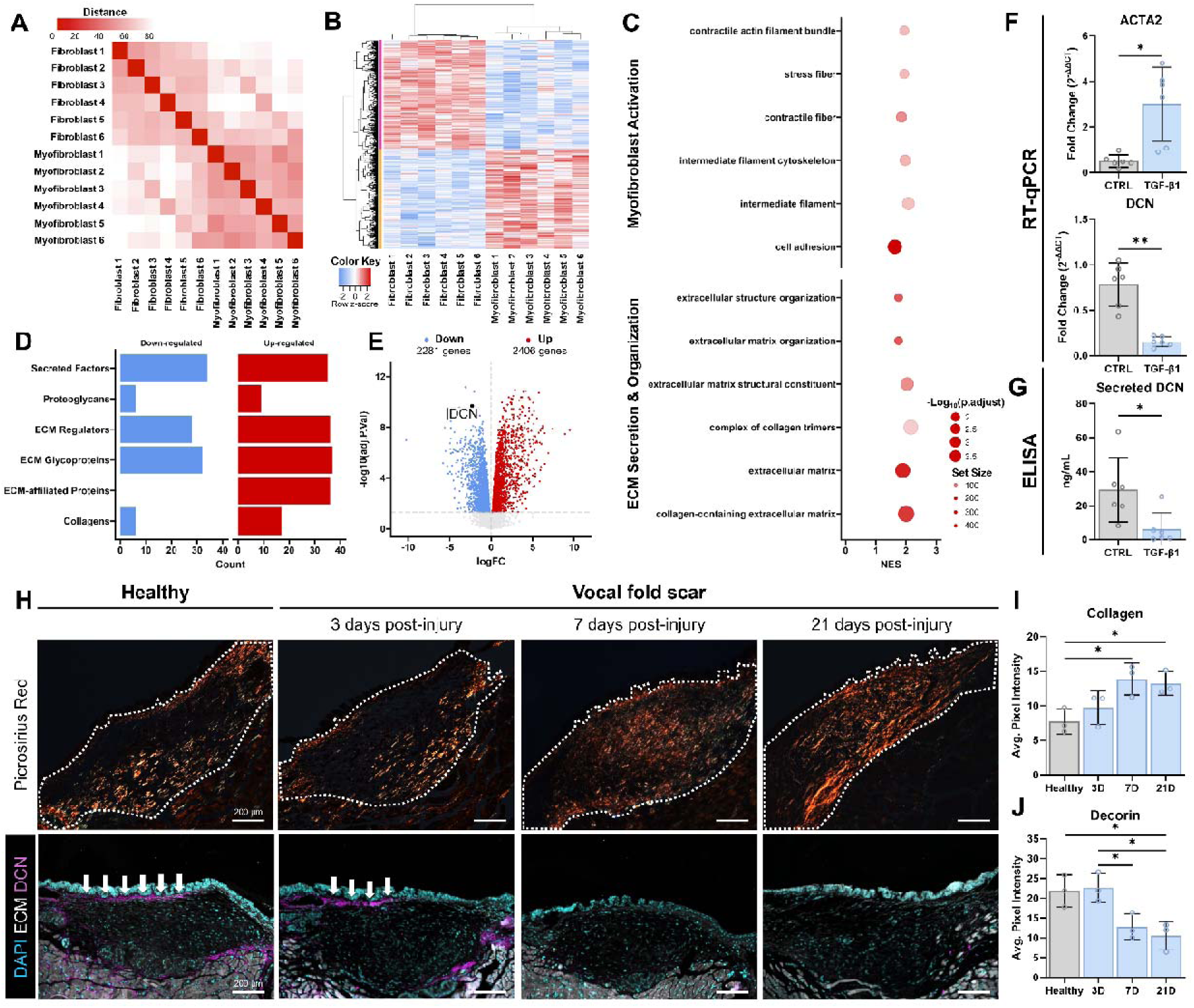
Decorin is downregulated in vocal fold myofibroblasts. (**A**) Distance matrix based on filtered counts per million of each gene visualized by heat map suggests that TGFβ1-induced vocal fold (VF) myofibroblasts have a unique transcriptional profile compared to resting VF fibroblasts. Darker red colors indicate more similarity between samples while lighter red and white colors indicate more dissimilarity between samples. (**B**) Heat map of differentially expressed genes (DEGs) shows differential clustering between TGFβ1-induced VF myofibroblasts and unstimulated VF fibroblasts. Red fill indicates increased gene expression in VF myofibroblasts, and blue fill indicates increased gene expression in VF fibroblasts. (**C**) Gene set enrichment analysis (GSEA) with Gene Ontology (GO) annotations indicates positive enrichment for pathways relating to myofibroblast activation and ECM secretion and organization in TGFβ1-stimulated cells. (**D**) Horizontal bar plot of DEGs and their matrisome annotation suggests significant changes to genes involved in extracellular matrix (ECM) secretion and maintenance. (**E**) Volcano plot of DEGs reveals that decorin (DCN) is among the most downregulated genes in VF myofibroblasts. Fold change values are based on gene expression in VF myofibroblasts relative to VF fibroblasts. (**F**) Reverse transcriptase-quantitative polymerase chain reaction (RT-qPCR) confirms TGF-β1 stimulation upregulates *ACTA2* and downregulates *DCN* in VF fibroblasts. (**G**) Secreted DCN is downregulated in TGFβ1-stimulated VF myofibroblasts as determined by enzyme linked immunosorbent assay (ELISA). (**H-J**) In a rat model of vocal fold scarring, collagen content increases while decorin content decreases *in vivo* in a time-dependent manner. *=p<0.05, **=p<0.01.

To translate these findings *in vivo,* we leveraged an endoscopy-guided abrasion injury model of VF scarring in rats (*29, 30*). Scarred VFs showed significant collagen accumulation at 7- and 21-days post-injury as determined by picrosirius red staining visualized by polarized light microscopy (Fig. 1H&I). DCN abundance was decreased during scar formation, reaching significance 21-days post-injury relative to healthy controls (Fig. 1H&J). Together, our *in vitro* results with human cells and *in vivo* findings using a rat model of VF scarring indicate that DCN is downregulated in scarred VFs, implicating DCN as important during disease progression.

### Decorin prevents TGFβ1-induced vocal fold myofibroblast activation

To demonstrate the antifibrotic capacity of DCN, cells were treated with or without exogenous DCN during TGF-β1 exposure (Fig. 2A). DCN treatment was sufficient to prevent TGFβ1-induced upregulation of the fibrosis-associated genes *COL1A1*, *COL5A1*, *ACTA2*, and *LOX* (Fig. 2B). DCN also prevented the incorporation of α-SMA into stress fibers (α-SMA^+^ stress fibers) (Fig. 2C, Fig. S2), one of the hallmarks of highly contractile myofibroblasts, which suggests reduced myofibroblast activation (*23*). Since substrate stiffness strongly impacts fibroblast phenotype *in vitro*, we next tested the antifibrotic properties of DCN in 3D methacrylated type I collagen hydrogels of a similar compressive modulus to native VF tissue, better mimicking the *in vivo* environment than a 2D plate (Fig. 2D). Cell-seeded gels exposed to TGF-β1 showed significant hydrogel contraction over 5 days, suggesting increased cell contractility following myofibroblast activation. However, constructs treated with both DCN and TGF-β1 showed minimal gel contraction (Fig. 2E) as well as reduced myofibroblast activation in 3D (Fig. S3). Histological analysis by hematoxylin & eosin (H&E) and by Masson’s trichrome staining indicate that DCN prevented TGFβ1-induced increases in overall ECM and collagen density (Fig. 2F).

**Fig. 2.**
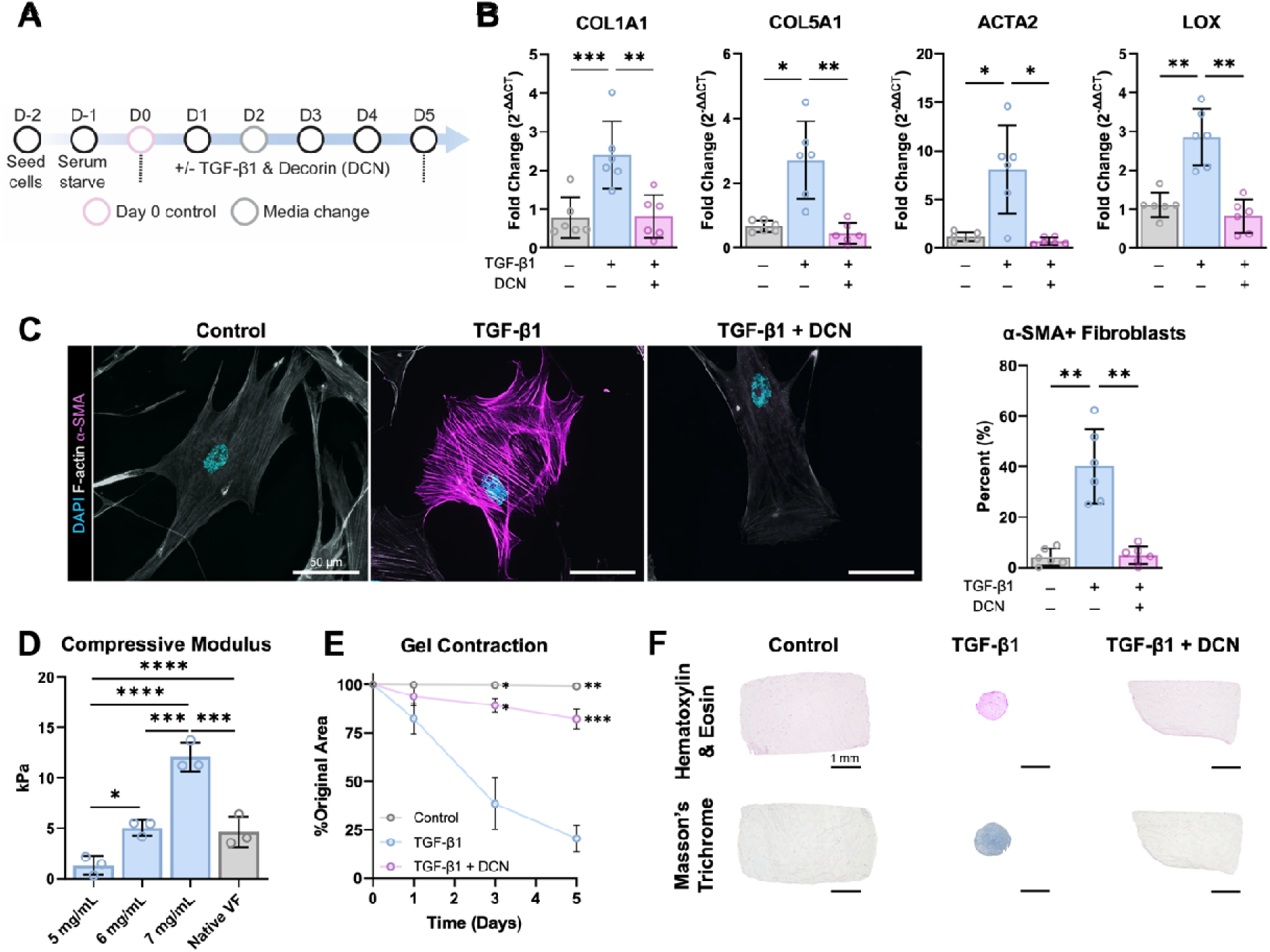
Exogenous decorin prevents TGFβ1-induced vocal fold myofibroblast activation *in vitro*. (**A**) Timeline of *in vitro* experiments. (**B**) Reverse transcriptase-quantitative polymerase chain reaction (RT-qPCR) shows that exogenous decorin (DCN) prevents TGFβ1-induced upregulation of pro-fibrotic *COL1A1, COL5A1, ACTA2,* and *LOX* gene expression. (**C**) Immunocytochemistry of the myofibroblast maker α-smooth muscle actin (α-SMA) indicates that TGF-β1 stimulation results in α-SMA incorporation into stress fibers, which can be prevented by DCN treatment. (**D**) Unconfined compression testing of methacrylated type I collagen hydrogels reveals concentration-dependent compressive moduli values. 6 mg/mL collagen are comparable to native VF tissue in terms of compressive modulus. (**E**) TGF-β1 treatment causes collagen hydrogel contraction in 6 mg/mL collagen hydrogels seeding with VF fibroblasts. DCN treatment reduces hydrogel contraction over 5 days. Significance represents comparisons to the TGF-β1 treated gels within the same time point. (**F**) Histological analysis via hematoxylin & eosin and Masson’s trichrome staining confirms that DCN treatment lessens TGFβ1-induced gel contraction and collagen secretion in collagen hydrogels. *=p<0.05, **=p<0.01, ***=p<0.005, ****=p<0.001.

### Hyaluronic acid microgels enable sustained decorin drug delivery

Direct injection of therapeutics into the VFs are limited by diffusion from the injection site, leading to a short residence time and a need for repeated injections, which are particularly impractical due to the complex upper airway anatomy (*16, 31*). Controlled drug delivery platforms that can be loaded with antifibrotic agents, such as DCN, are then necessary to achieve therapeutic potential and reduce injection frequency. To this end, we engineered hyaluronic acid microgels by a modified extrusion fragmentation method including sonication and homogenization steps (Fig. 3A&B). Hyaluronic acid was selected for its ability to withstand the small amplitude, high frequency tissue oscillations during phonation and for its long-standing use as a IML material, providing physicians with a sense of familiarity and a history of safe use. Microgels were engineered to achieve a mean microgel diameter of 26.48 μm (95% CI: 24.00, 28.96) (Fig. 3C). We observed no significant differences in mean microgel diameter between batches, suggesting low batch-to-batch variability (Fig. 3C). Since intracordal injections are performed with a 26 gauge (G) syringe needle, we ensured that extrusion through a 26G needle did not lead to additional microgel fragmentation or significantly affect microgel size (Fig. 3C, Fig. S4). Additionally, we show that microgel extrusion through a 26G needle requires sub-kilonewton forces and is comparable to a clinically-used IML material, Prolaryn® Plus, which is an injectable hydroxyapatite microsphere-based system. We also determined that microgels with an average diameter of 70.10 μm and 109.01 μm, respectively, are easily injected with sub-kilonewton forces (Fig. 3D), demonstrating the tunability of this platform.

**Fig. 3.**
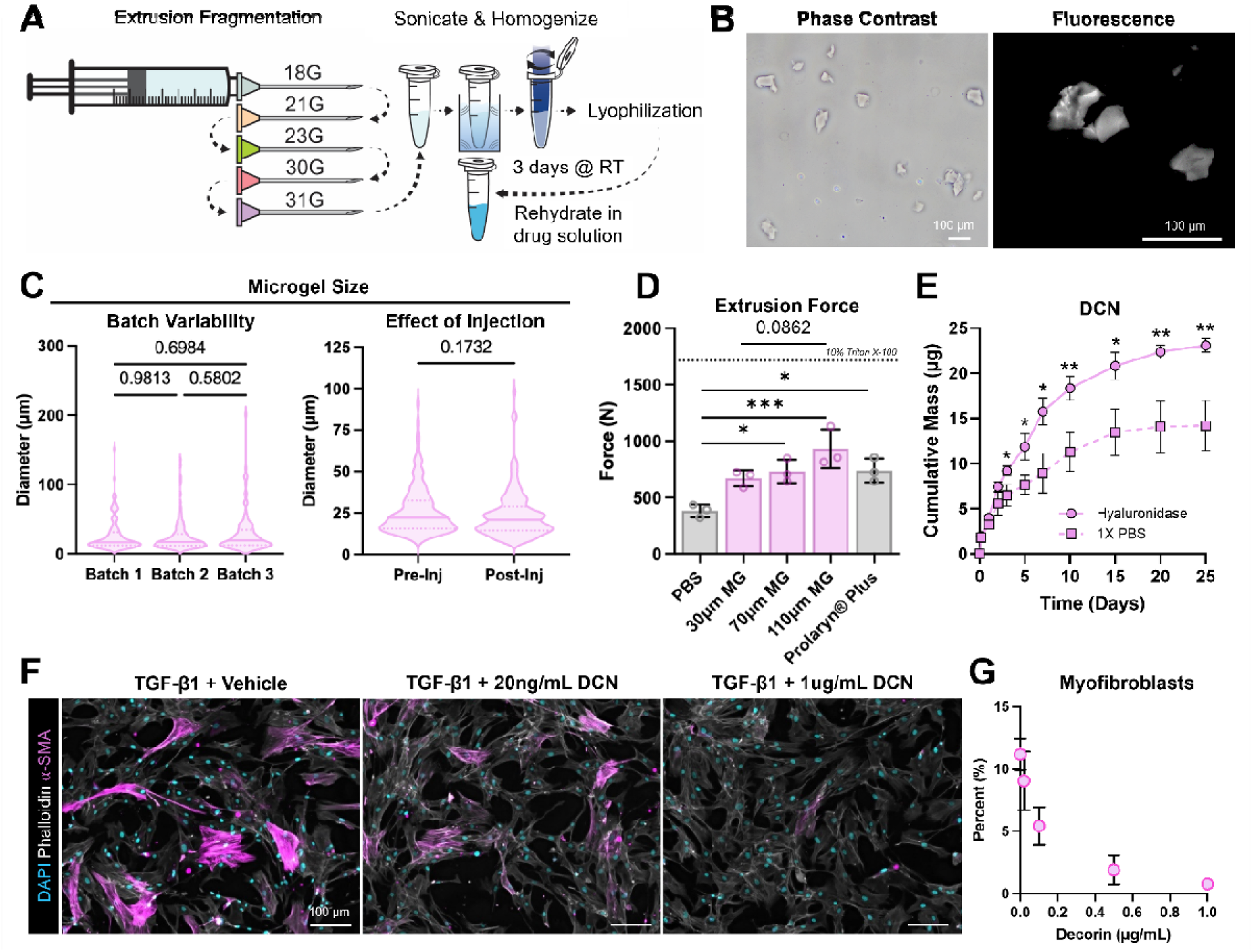
Hyaluronic acid microgels enable sustained release of decorin. (**A**) Schematic of hyaluronic acid microgel fabrication process via a modified extrusion fragmentation protocol. Bulk hyaluronic acid hydrogel is formed via photopolymerization and loaded into a syringe. Material is consecutively extruded through needles with smaller diameters, sonicated, homogenized, and then lyophilized. Decorin (DCN) is loaded into the material via passive loading. (**B**) Representative phase-contrast optical microscopy and fluorescence images of FITC-dextran loaded microgels. (**C**) Microgels show low batch-to-batch variability in microgel size and no significant differences in pre- and post-injection using a 27G needle, which is within the range of needle sizes used for *in vivo* applications. (**D**) Using a force sensor attached to a syringe with a 26G needle, extrusion force was determined for PBS, our engineered microgels of varying sizes, and Prolaryn® Plus (clinical standard), and. Our platform required a similar sub-kilonewton extrusion force to a clinically-used IML material. (**E**) Hyaluronic acid microgels show controlled release of DCN over 25 days in both 1X PBS and a hyaluronidase solution. Significance represents the comparison between released DCN in hyaluronidase-containing solution and 1X PBS. (**F&G**) DCN releasate retains antifibrotic properties as indicated by reduced α-SMA expression in vocal fold (VF) fibroblasts treated with TGF-β1. *=p<0.05

For passive drug loading, lyophilized microgels were rehydrated in DI water containing DCN (Fig. 3A). Since hyaluronic acid, and thus our microgel platform, can be degraded by endogenous hyaluronidases after intracordal injection (*32*), we evaluated the DCN release profile at 37 in both PBS and a hyaluronidase solution. In both conditions, DCN-loaded microgels exhibited an initial burst release followed by a near-linear, sustained drug release profile (Fig. 3E). In PBS, this near-linear region continued for approximately 2 weeks, at which point drug release began to plateau. Beginning at 3 days, cumulative DCN release was significantly higher in a hyaluronidase solution compared to PBS (Fig. 3E). The near-linear region lasted for 1 week in the hyaluronidase solution followed by sustained drug release for a total of 25 days (Fig. 3E). Releasate from DCN-loaded microgels was collected then used to treat VF fibroblasts during TGF-β1 exposure. Microgel releasate significantly reduced the percentage of VF fibroblasts with α-SMA-positive stress fibers, suggesting that DCN retained its bioactivity after microgel encapsulation and that DCN-loaded microgels could reduce myofibroblast activation (Fig. 3F&G). Together, these data demonstrate that hyaluronic acid microgels enable controlled DCN drug delivery with retained protein bioactivity.

### Decorin-loaded microgels prevent vocal fold scarring in a rat model

Given the narrow airway lumen and small size of VFs, intracordal injections in small rodent animal models are non-trivial. To visualize and inject rat VFs, we developed and implemented a transoral endoscopy-guided approach, which we then combined with our existing model of VF scarring to test the feasibility and efficacy of DCN-loaded microgels *in vivo* (Fig. 4A, Movie S1). VFs were visualized under direct laryngeal endoscopy and were then left untreated or injected with either blank microgels or DCN-loaded microgels using a 27G needle. Microgels were successfully injected into the thyroarytenoid muscle (vocalis muscle), which is immediately lateral to the VF lamina propria, in rats (Fig. 4B&C, Fig. S5). After intracordal injection of microgels, rats were then subjected to an endoscopy-guided bilateral abrasion injury using a wire brush to induce VF scarring.

**Fig. 4.**
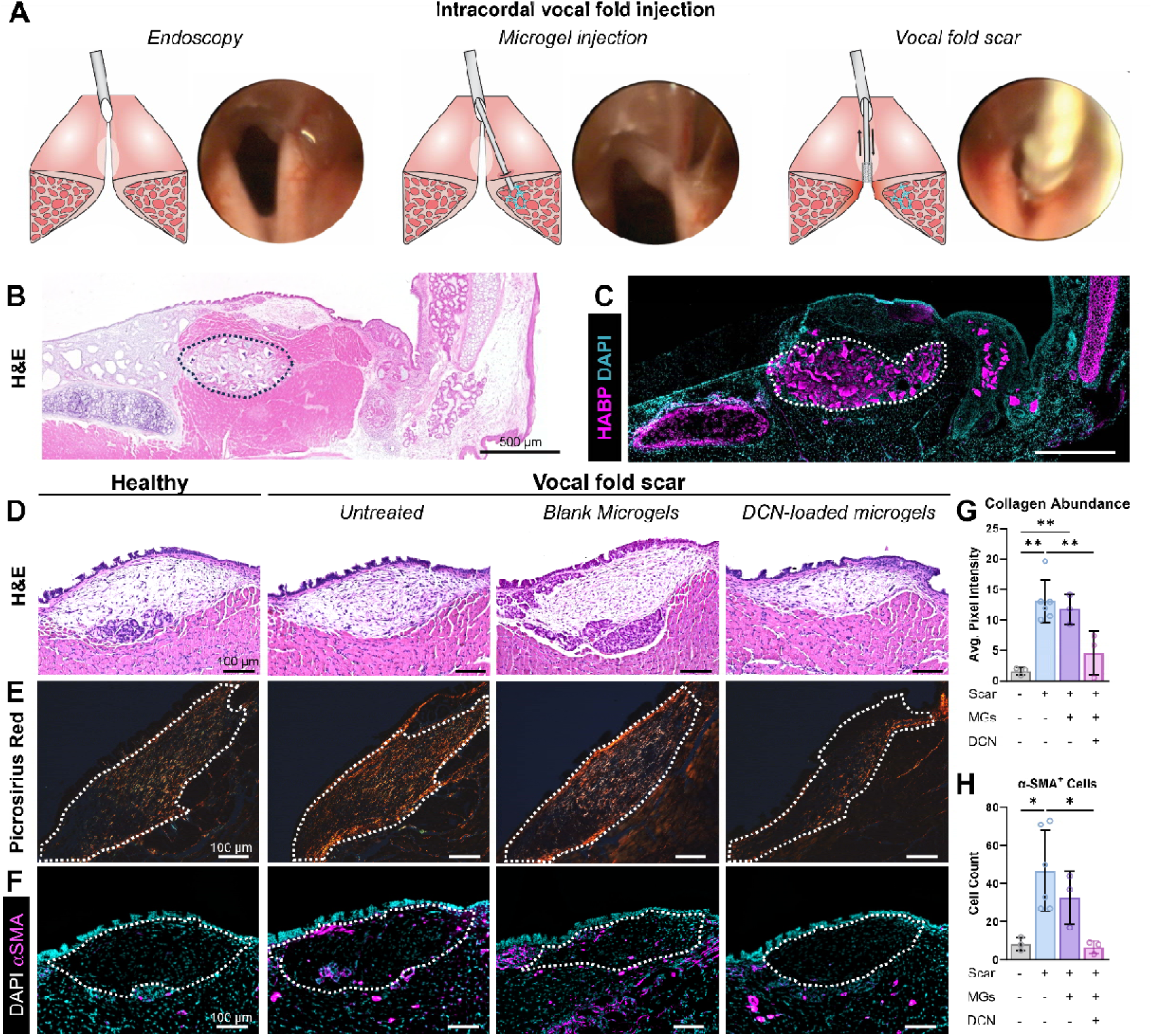
Decorin-loaded hyaluronic acid microgels prevent vocal fold scarring in a rat model. (**A**) Schematic and endoscopy images of endoscopy-guided intracordal injection and vocal fold (VF) scarring rat model. (**B**) Hematoxylin & eosin (H&E) and (**C**) hyaluronan binding protein (HABP) histology of rat larynges injected with blank hyaluronic acid MGs show successful intracordal injection of the underlying vocalis muscle. Dotted outlines are drawn around injected microgels. (**D**) H&E staining of mid-coronal vocal fold sections suggests a denser extracellular matrix in scarred VFs left untreated or injected with blank microgels relative to healthy rats and injured rats injected with DCN-loaded microgels at 7 days post-injury. (**E&G**) Picrosirius red staining visualized by polarized light microscopy show that VF scarring is characterized by significant collagen accumulation, which is unaffected by injection of blank microgels. DCN-loaded microgels were sufficient to prevent collagen accumulation. Dotted outlines drawn around the VF lamina propria for each representative image. (**F&H**) DCN-MGs mitigates persistent myofibroblast activation *in vivo* as evidenced by reduced α-SMA (myofibroblast marker) immunostaining *in* vivo. Dotted outlines drawn around the VF lamina propria for each representative image. *=p<0.05, **=p<0.01

Mechanical VF injury resulted in higher ECM density in rats receiving either no treatment or blank microgel injections relative to healthy controls. However, DCN-loaded microgels prevented injury-induced increases in ECM density, and rats in this experimental group presented similar to uninjured animals (Fig. 4D). We performed picrosirius red staining and visualized collagen fibers by polarized light microscopy to evaluate collagen accumulation after VF injury. Animals receiving either no treatment or a blank microgel injection exhibited significant collagen accumulation 7 days post-injury (Fig. 4E&G). However, rats injected with DCN-loaded microgels showed no significant collagen accumulation post-injury, suggesting scarless tissue repair (Fig. 4E&G). Since myofibroblasts are the key orchestrators of scar formation, we performed immunostaining for the classical myofibroblast marker α-SMA. As expected, healthy, uninjured rat VFs contain few myofibroblasts with low α-SMA expression (Fig. 4F&H). However, mechanical VF injury induced significant myofibroblast activation, which was not strongly affected by blank microgel injections (Fig. 4F&H). On the other hand, DCN-loaded microgels administrated via intracordal injection reduced myofibroblast activation at 7-days post-injury (Fig. 4F&H), which indicates physiologic rather than fibrotic wound healing. Together, these data establish the feasibility of DCN-loaded microgels as an antifibrotic treatment for VF scarring after direct airway injury.

To demonstrate clinical applicability of this platform, we also injected our engineered microgels into the VF lamina propria in a pig model. Microgel injection resulted in VF medialization via endoscopy, and histological analysis confirmed successful intracordal injection (Fig. S6).

## Discussion

VF scarring remains a significant clinical challenge and is the leading refractory cause of poor voice after VF injury (*8, 9*). Upon injury, immune cells infiltrate the wound bed and secrete a complex array of signaling molecules, including numerous proinflammatory cytokines (*33–35*). These inflammatory mediators initiate key signaling cascades in resident VF fibroblasts, inducing myofibroblast activation. VF myofibroblasts play a central role in wound healing because they are responsible for wound closure, ECM deposition, and tissue remodeling (*36*). Scar formation begins upon prolonged myofibroblast activation and persistence, which leads to wound contraction, excessive matrix accumulation, and a disorganized ECM architecture. Fibrotic changes to the structure of the VF lamina propria disrupt biomechanical properties and, as a result, patients with VF scarring experience severe dysphonia (*37*) and significant patient morbidity (*2, 38*).

There are currently no successful non-surgical treatments for VF scarring with consistent, robust positive outcomes. The two primary clinical interventions are intracordal steroid injections and IML. In addition to their low clinical efficacy, steroid injections are associated with adverse events such as atrophy of the VFs, mucosal glands, or vocalis muscle, especially after repeated administration (*20, 39*). Safety concerns are heightened in pediatric populations, further emphasizing the limited utility of this approach. IML is performed on patients with glottic incompetence to medialize the VFs and improve phonatory outcomes (*13*). While temporarily successful, IML is not disease-modifying and fails to target any mechanisms driving scar formation.

In this work, we first performed transcriptomics of human VF fibroblasts and TGFβ1-induced VF myofibroblasts to uncover differentially expressed genes (DEGs) that may be critical for VF myofibroblast activation and function. Our RNA sequencing revealed that DCN is among the most downregulated genes in the transcriptome of TGFβ1-induced VF myofibroblasts (Fig. 1E). We also observed decreased intracellular and secreted DCN protein in VF myofibroblasts (Fig. S1B, Fig. 1G), in support of our transcriptomics data. Consistent with previous reports (*21, 40*), we found that DCN is abundant and concentrated to the superficial layer of the VF lamina propria in healthy animals and at the earliest stages of wound healing. We also demonstrate that DCN rapidly disappears as fibrotic wound healing continues and a mature scar forms (Fig. 1H&J). These data are consistent with those from Ohno et al. showing decreased DCN gene expression in injured rat VFs (*41*) as well as with the results from Thibeault et al. who identified decreased DCN expression in injured rabbit VFs (*42*). Logically, we then sought to determine whether restoring DCN presentation improved scarless healing by modulating fibroblast behavior.

During TGF-β1 stimulation, we treated VF fibroblasts with DCN to evaluate its antifibrotic capacity and found that DCN treatment prevented TGFβ1-induced upregulation of the pro-fibrotic genes *COL1A1*, *COL1A2*, *ACTA2*, and *LOX* (Fig. 2B) as well as formation of α-SMA^+^ stress fibers (Fig. 2C). We tested these findings in a more biomimetic 3D environment by embedding VF fibroblasts in collagen hydrogels with similar compressive moduli to native VFs (Fig. 2D). Because upper airway myofibroblasts are highly contractile (*24, 43*), TGF-β1 treatment resulted in collagen hydrogel contraction over 5 days; however, DCN presentation significantly decreased gel contraction, pointing to a reduction in myofibroblast activation (Fig. 2E). Our findings are consistent with work from Zhang et al. who demonstrate that DCN inhibits collagen gel contraction by both normal and hypertrophic scar fibroblasts (*44*). These data suggest that DCN is sufficient to prevent myofibroblast activation in human VF fibroblasts *in vitro*.

DCN is a key signaling molecule and an important structural ECM component, both of which may confer anti-scarring benefits. Biochemically, DCN has an extensive and diverse interactome (*45, 46*), and it is likely that DCN modulates myofibroblast activation through multiple mechanisms acting in concert. DCN has known binding sites for a number of proinflammatory or profibrotic molecules, including TGF-β1, tumor necrosis factor-α (TNF-α), cellular communication network factor 2 (CCN2), and WNT1-induced signaling pathway protein 1 (WISP1) (*45, 46*), all of which have been implicated in scar formation across organ systems. DCN also binds multiple receptor tyrosine kinases (RTKs) with high affinity, functioning as a pan-RTK inhibitor (*47*). RTK inhibition may offer one possible explanation for DCN’s antifibrotic effects, supported by the use of the FDA-approved pan-RTK inhibitor nintedanib as a treatment for idiopathic pulmonary fibrosis (*48*). Since TGF-β signaling is central to VF myofibroblast activation and scar formation (*10*), DCN’s ability to sequester TGF-β1 and diminish TGF-β signaling likely gives rise, in part, to its antifibrotic potential. Anti-scarring approaches aimed at reducing TGF-β activity, such as through small interfering RNA (siRNA) targeting *Smad3*, one of the predominant effector proteins in canonical TGF-β signaling, have shown promise across different models of VF scarring (*49, 50*). In a swine model of airway intubation injury, Miar *et al.* demonstrated that controlled delivery of *Smad3-*targeting siRNA limits myofibroblast activation and reduces tissue stiffness (*51*). Furthermore, as a structural ECM protein, DCN also binds to collagen fibers (*52, 53*) and acts as a regulator of collagen fibrillogenesis and collagen network assembly (*46, 54*), both of which are dysregulated in VF scarring. Across various tissue types such as tendon (*55*), skin (*56*), and cornea (*57*), DCN-null mice present with disorganized and irregularly shaped collagen fibers, disrupting tissue biomechanics. Exogenous DCN treatment may improve functional outcomes in VF scarring by simultaneously reducing pro-fibrotic cell signaling and restoring native ECM collagen fibrillogenesis. Nonetheless, additional studies are required to dissect the relative contribution of each DCN binding partner and the structural effects of DCN in the context of VF scarring.

To minimize the risk of off-target effects, reduce drug diffusion from the injection site, and improve the therapeutic efficacy of DCN, we engineered microgels for controlled drug delivery to the VFs (Fig. 3A). Hyaluronic acid was selected for microgel fabrication because it is abundant in the VF lamina propria throughout tissue maturation (*21*); furthermore, hyaluronic acid-based biomaterials are often injected in the VF for IML (*58*). Since hyaluronic acid can dampen vibrational forces and act as a shock absorber, its use helps to lessen the risk of VF injuries during phonation, where constant vibrations induce complex and continuous stresses that may induce chronic VF damage (*59*). We fabricated microgels with a mean diameter of 23.53 μm post-injection (Fig. 3C), achieving a size distribution similar to that of another clinically-used microparticle-based IML biomaterial, Prolaryn® Plus, which contains hydroxyapatite microspheres between 25−45 μm in diameter. Our microgels are easily injected with sub-kilonewton forces through a 26G needle (Fig. 3D), requiring a lower extrusion force than other hyaluronic acid-based biomaterials frequently administered via intracordal injection (*59–61*). DCN is readily loaded into hyaluronic acid microgels for controlled drug delivery over 25 days (Fig. 3E). This is an ideal timeline since the transition from fibrogenesis to matrix remodeling occurs during 3 4 weeks in rats (*62*).

We established the feasibility of DCN-loaded hyaluronic acid microgels by first performing successful injections in both rats (Fig. 4B&C) and then in pigs, whose VFs are closer in size to humans (Fig. S6). Then, we demonstrated anti-fibrotic efficacy in a preclinical rat model of VF scarring (Fig. 4A−C). As expected, VF injury by abrasion with a wire brush resulted in increased collagen accumulation and significant myofibroblast activation at 7 days post-injury. In agreement with Catten et al., myofibroblasts were most abundant in the superficial layer of the VF lamina propria (*22*). Blank microgel injection showed no significant effect on fibrotic wound healing, reinforcing the need for the delivery of an antifibrotic such as DCN. In fact, DCN-loaded microgels prevented all measured outcomes of VF scarring in our system, decreasing collagen accumulation and reducing myofibroblast activation (Fig. 4D−H). Our results are consistent with the observations from several other groups studying cardiac and corneal fibrosis in that DCN has antifibrotic properties. Mohindra et al. leveraged an ischemia reperfusion myocardial infarction rat model to demonstrate the controlled delivery of DCN from hydrogel microrods attenuated cardiac fibrosis, with decreased collagen accumulation and improved cardiac performance (*63*). Similarly, Hill et al. showed that sustained release of DCN from a gellan-based fluid gel eye drop system promoted scarless corneal tissue repair in a microbial keratitis murine model (*64*). Altogether, our study provides proof-of-concept feasibility for DCN-loaded hyaluronic acid microgels as a novel antifibrotic VF scarring therapy.

In this work, we show that the proteoglycan decorin (DCN), which is abundant in healthy VFs, is downregulated in VF myofibroblasts and depleted in scarred VFs. We demonstrate that the exogenous administration of DCN prevents TGFβ1-induced myofibroblast activation in human VF fibroblasts, evidenced by reduced pro-fibrotic gene expression, α-SMA^+^ stress fibers, and cell contractility. To overcome drug delivery challenges after intracordal injection, we loaded DCN into hyaluronic acid microgels fabricated via extrusion fragmentation. Microgels demonstrated sustained DCN delivery for over 3 4 weeks with retained protein bioactivity. In a preclinical rat model of vocal fold scarring, DCN-loaded microgels reduced collagen deposition and myofibroblast activation compared to both untreated and blank microgel treated injuries, restoring a tissue phenotype consistent with scarless healing. These findings highlight the therapeutic potential of this platform to prevent fibrotic remodeling and improve outcomes in vocal fold repair.

## Materials and Methods

### Materials and Reagents

Dulbecco’s modified eagle medium (DMEM), fetal bovine serum (FBS), Gibco minimum essential medium (MEM) non-essential amino acids, Gibco Geneticin™ selective antibiotic (G418 Sulfate), TrypLE™ express enzyme, human TGF-β1 recombinant protein, acetic acid, 1X PBS, 10X PBS, SuperScript™ IV first-strand synthesis system, PowerUp™ SYBR™ green master nix, 10% buffered formalin, xylene, Epredia™ precision cut paraffin, normal goat serum, bovine serum albumin (BSA) DNase- and protease-free powder, Epredia™ harris hematoxylin non-acidified, Epredia™ eosin-Y alcoholic, Fluoromount-G™ mounting medium, and NucBlue™ Live ReadyProbes™ reagent (Hoechst 33342) were purchased from ThermoFisher Scientific (Waltham, MA). Penicillin-streptomycin (P/S), lithium phenyl-2,4,6-trimethylbenzophosphinate (LAP), sodium hydroxide (NaOH), Triton X-100, and hyaluronidase from bovine testes were purchased from MilliporeSigma (Burlington, MA). Citric acid trisodium salt dihydrate was purchased from VWR (Radnor, PA). Picrosirius red was purchased from StatLab (McKinney, TX). Human TGF-β1 DuoSet ELISA was purchased from R&D Systems (Minneapolis, MN) and used following manufacturer instructions. Human his-tag decorin protein was purchased from Acro Biosystems (Newark, DE). RNeasy® plus mini kit and proteinase K was purchased from Qiagen (Hilden, Germany) and used following manufacturer instructions. All primers used for RT-qPCR were purchased via Integrated DNA Technologies (Coralville, IA). Methacrylate type I bovine collagen and methacrylated hyaluronic acid was purchased from Advanced Biomatrix (Gothenburg, Sweden).

### Cell Culture

Human vocal fold fibroblasts were provided as a kind gift from the laboratory of Dr. Susan Thibeault, whose team immortalized vocal fold fibroblasts isolated from deceased individuals showing no signs of vocal fold damage (*24, 65, 66*). Patient information can be found in our previous publication (*12*). Human vocal fold fibroblasts were cultured in a monolayer on tissue-culture plastic at 37°C with 5% CO_2_ with DMEM supplemented with 10% FBS, 2% penicillin-streptomycin, 1% MEM NEAA, and 200 μg/ml G418 sulfate. Cells were seeded onto 6-well plates (#215-105, GenClone) at a seeding density of 50,000 cells/well or onto 8-well removable frames on lumox® slides (#94.6150.801, Sarstedt) at a seeding density of 7,000 cells/well, unless otherwise specified. Media changes were performed every 2-3 days for the course of all *in vitro* experiments, unless otherwise specified.

To induce myofibroblast activation on vocal fold fibroblasts grown atop tissue-culture plastic, cells were stimulated with DMEM supplemented with 10ng/mL of human TGF-β1 recombinant protein (#100-21, ThermoFisher Scientific), 2.5% FBS, 2% penicillin-streptomycin, and 1% MEM NEAA for 5 days. Exogenous decorin stimulation was achieved by also stimulating with DMEM supplemented with 10ug/mL human his-tag decorin protein (#DE1-H5223, Acro Biosystems).

### RNA isolation and bulk RNA Sequencing

Please refer to our group’s previous work describing our full RNA isolation and bulk RNA sequencing pipeline (*12*). In short, total RNA was extracted using the Maxwell RSC simply RNA kit (#AS1390, Promega) in accordance with manufacturer instructions. Nanodrop was used to verify RNA concentration, yield, and purity. RNA integrity value (RIN) was determined by RNA TapeStation 4200 (Agilent). Illumina Stranded Poly-A Selection RNA sequencing kit was performed by the Children’s Hospital of Philadelphia Center for Applied Genomics. After preparation with 100 ng of starting input, samples were sequenced on a S2 200 cycle kit on a NovaSeq 6000. Output per-cycle binary base call (BCL) files from Illumina sequencing instruments were demultiplexed with the Illumina DRAGEN-bioIT platform using the bcl2fastq program. Sample specific FASTQ files were then filtered to remove adaptor sequences, low-quality tags with unknown nucleotides greater than 10%, and reads with greater than 50% low quality bases. All reads that passed quality control parameters were mapped using STAR 2.7.3a-GCC-9.30 using the GRCh38 genome.

All transcriptome analysis was carried out in with R (version v4.3.0) in RStudio (version 2021.09.0+351) with R packages collected from Bioconductor. Please refer to Table S2 for a comprehensive list of all software used in this work. Normalized counts per million (CPM) were used for gene expression analysis through methods provided in EdgeR (version 3.42.2). Heatmaps of sample-to-sample distances as well as of gene expression were created, and differentially expressed genes were determined using the limma package (version 3.56.1). Next, gene set enrichment analysis was performed with Gene Ontology (GO) annotations using clusterProfiler (version 4.8.1) to evaluate gene sets with positive or negative enrichment were determined.

### RNA isolation and RT-qPCR

RNA was extracted and purified using the RNeasy Plus Mini Kit (#74134, Qiagen) following manufacturer instructions. RNA quantification was quantified and then converted to cDNA via reverse transcription with SuperScript IV First-Strand Synthesis System kit (#18091050, Invitrogen) and random hexamer primers. Real-time quantitative reverse transcription polymerase chain reaction (RT-qPCR) was performed with a MiniAmp™ Thermal Cycler (Applied Biosystems). 2ng of cDNA, 0.5μM of forward primers, 0.5μM of reverse primers, and PowerUp™ SYBR™ Green Master Mix (#A25742, Applied Biosystems) was mixed in a 384-well plate for RT-qPCR. Run time was as follows: 1 cycle of 50°C for 2 min and 1 cycle of 95°C for 2 min; 40 cycles of 95°C for 1 second and 60°C for 30 seconds; 1 cycle of 95°C for 15 seconds, 1 cycle of 60°C for 1 minute, and 95°C for 15 seconds. Gene expression was evaluated as fold change, which was calculated as 2^−ΔΔCT^ using comparative cycle threshold in reference to a day 0 control. Please refer to Table S3 for a description of all primers used for RT-qPCR.

### Collagen hydrogel fabrication, characterization, and cell seeding

Methacrylate type I bovine collagen (#5270, Advanced Biomatrix) was resuspended to 8mg/mL per manufactuer instructions in 20mM acetic acid. Collagen was diluted to achieve a final concentration of 5, 6, or 7 mg/mL using 1M NaOH, 10X PBS, and lithium phenyl-2,4,6-trimethylbenzophosphinate (LAP). For cell experiments, vocal fold fibroblasts were added with supplemented DMEM to achieve a seeding density of 500,00 cells scaffold. Components were well-mixed prior to pipetting into sterilized silicone molds and subsequent incubation at 37°C for 2 h. Molds were then removed without disrupting gel integrity. Acellular gels were stored in PBS until mechanical testing and cellular gels were suspended in supplemented DMEM and cultured under agitation at 100 rpm. After 24 h, the cell-laden constructs were treated with or without TGF-β1 and exogenous decorin (DCN). Images were acquired over 5 days to monitor collagen hydrogel contraction. Collagen hydrogel area was measured with ImageJ software.

### Histology and immunofluorescence

For excised laryngotracheal complexes and cells embedded in collagen gels, samples were fixed in 10% buffered formalin overnight at 4. Samples were washed in 1X PBS, dehydrated via an ethanol gradient diluted in 1X PBS, cleared with xylene washes, and then embedded in paraffin. 8 μm-thick coronal paraffin-embedded sections were collected using a Microm HM 355S (Thermofisher Scientific) and stored at room temperature after drying overnight. For histology, sections were deparaffinized, rehydrated and routinely stained with hematoxylin & eosin or picrosirius red prior to visualization with bright-field microscopy or polarized-light microscopy. For standard immunofluorescence, samples were deparaffinization and rehydrated. Antigen retrieval was then performed by incubation in either proteinase K diluted in 1X PBS at room temperature for 30 min or in sodium citrate (pH 6) buffer at 60°C for 2 hours. For immunofluorescence of cells cultured in tissue-culture plastic, samples were fixed with 10% buffered formalin for 15 min at room temperature followed by 1X PBS washes. All samples were incubated in blocking buffer (0.1% Triton-X 100, 10% goat serum, and 2% BSA) for 2 h at room temperature prior to overnight primary antibody incubation. Following 1X PBS washes, secondary antibodies were applied for 1 h and 30 min at room temperature with DAPI (Table S4). Samples were mounted using Fluromount-G. Bright-field and fluorescence microscopy images were acquired using a BZ-X810 fluorescence microscopy (Keyence). Polarized light microscopy was performed for picrosirius red-stained sections using a MicorPublisher 6 microscope camera (Teledyne). All images were analyzed using ImageJ software.

### Microgel fabrication and characterization

Methacrylated hyaluronic acid was purchased (#5274, Advanced Biomatrix), and bulk hyaluronic acid hydrogels were fabricated via photopolymerization with LAP as photoinitator following manufacturer instructions. Bulk hyaluronic acid hydrogel was extruded through different sized needles, including 18G, 21G, 23G, and 30G to create larger fragmented microgels. For medium and small microgels, material was also extruded through a 31G needle. Deionized water was flushed through syringe needles after extrusion to clear the needles of residual material. To create small microgels, samples were then centrifuged at 3000g for 5 minutes, homogenized for 1-1.5 minutes, vortexed for 5 minutes, and then sonicated at for 2 minutes. Gel pellets were centrifuged at 6000g for 5 minutes, frozen at −80 m and then lyophilized. After lyophilization, samples were stored at −20 until rehydration. Microgel size was determined via bright-field microscopy (Eclipse Ts2, Nikon) with phase-contrast followed by analysis using ImageJ software.

Extrusion force measurements were collected on PBS, Prolaryn® Plus, and our microgel platform with a custom Arduino-based force measurement setup, similar to previously reported methods (*67, 68*). In short, the force required to extrude a sample of material through a 27G syringe needle was determined with a force-resistive sensor. A 1mL syringe was loaded with the material of interest and the force sensor was placed between the syringe plunger and the pusher block of a syringe pump. 0.5mL of material was extruded at 200μL/min and arbitrary output voltage values were collected over time. These results were calibrated to force measurements through a calibration curve collected by placing known weights on the force sensor. After generating a force versus time curve, the average of forces in the plateau region of the force-time curve was determined.

To determine the release profile of DCN from this platform, we used 1mL of methacrylated hyaluronic acid pre-cursor to fabricate hyaluronic acid microgels. Samples were rehydrated in either deionized water or 15 μg/mL human his-tag decorin protein (#DE1-H5223, Acro Biosystems) in deionized water for 3 days at room temperature. Release profiles were determined for DCN microgels in PBS and 5U hyaluronidase (#H3506, Sigma-Aldrich) by incubating the samples at 37 at 100rpm. The bioactivity of released protein was confirmed by collecting the releasate from DCN-loaded microgels after 3 days. Vocal fold fibroblasts were treated with TGF-β1 supplemented with or without DCN-loaded microgel releasate to achieve a final concentration of 0-1μg/mL of DCN. Cells were cultured for 5 days prior to immunocytochemistry for α-SMA and subsequent quantification using ImageJ software. The percentage of cells positive for α-SMA was determined by dividing the number of cells with α-SMA staining on cell stress fibers by the total number of nuclei, which was counted using the built-in analyze particles analysis tool.

### Animal Studies

Male wild-type Sprague Dawley rats of 15-16 weeks of age were used following IAC 22-001462 at the Children’s Hospital of Philadelphia Institutional Animal Care and Use Committee. During the study, rats were housed in a temperature-controlled specific-pathogen-free facility. Rats were anesthetized by isoflurane, and low-dose buprenorphine SR (0.6-1.2 mg/kg) was administered via subcutaneous injection. Under continuous isoflurane, VFs were visualized under direct laryngeal endoscopy with a 1.9 mm-diameter 30° angled endoscope (#R64301BA, Karl Storz) with an operating sheath (#61029D, Karl Storz). Some rats received intracordal injection prior to VF injury using a custom 27G, 6.5-inch, point style 4 needle (#7803-01, Hamilton Company) attached to a removable needle 25-microliter syringe (#80430, Hamilton Company) passed through the operating sheath. VF scarring was then induced by endoscopy-guided brushing the airway 10-15 times using a custom micro-spiral stainless teel brush (#460010, Gordon Brush). Animals were then housed as described above until experimental endpoint at either 3-, 7-, or 21-days post-injury.

### Statistical Analysis

Statistical analysis was performed and data were plotted in Prism GraphPad 9.0. All data were graphed and reported as means ± standard deviation. For fibroblast *in vitro* assays, paired t tests or one-way analysis of variance (ANOVA) with Tukey’s multiple comparisons was used to determine statistically significant differences. Histological analyses were performed with one-way ANOVA with Tukey’s multiple comparisons was used to determine statistically significant differences between treatment conditions. P < 0.05 is considered significant.

## Acknowledgments

We thank the laboratory of Véronique Lefebvre, Ph.D for their histological facilities.

## Funding

Children’s Hospital of Philadelphia Research Institute (RG, KZ)

Frontier Program in Airway Disorders of the Children’s Hospital of Philadelphia (RG)

Ri.MED Foundation (RG)

National Science Foundation Graduate Research Fellowship Program No. DGE 1845298 (RMF, MRA)

NIH NIAMS P30AR069619 for Core Facility use

## Author contributions

Conceptualization: RMF, KBZ, RG

Methodology: RMF, EAB, KSM, KBZ

Software: RMF, MRA

Investigation: RMF, EAB, EAB, HMO, HB, MRA, YG

Visualization: RMF

Writing—original draft: RMF, RG

Writing—review & editing: RMF, KBZ, RG

Supervision: RMF, RG

Funding Acquisition—RMF, KBZ, RG

## Supplementary Materials

**Fig. S1.**
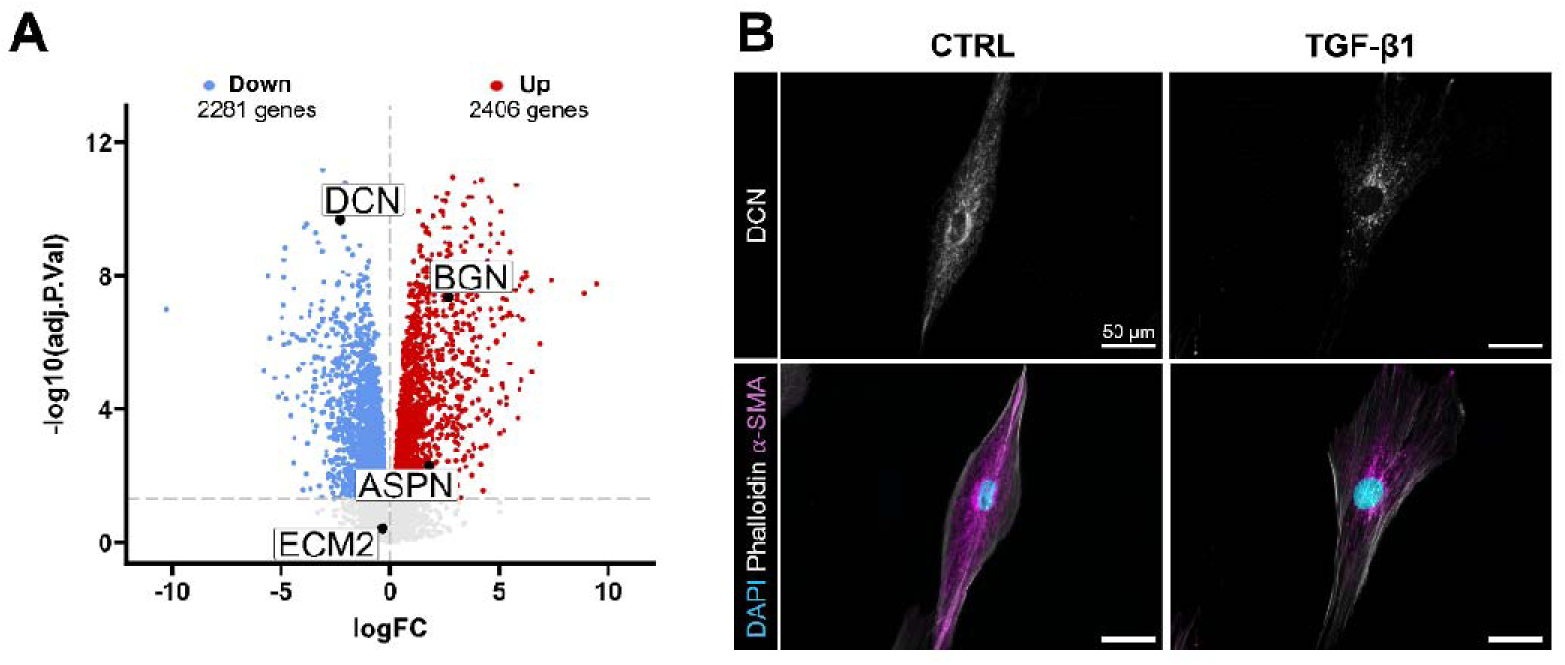
VF myofibroblasts show reduced decorin gene expression and intracellular protein abundance. (A) Volcano plot of bulk RNA sequencing experiments comparing differentially expressed genes in VF myofibroblasts to resting VF fibroblast controls. DCN is among the most downregulated genes, unlike other small leucine-rich proteoglycans of the same family and class. (B) Immunocytochemistry of DCN in human VF fibroblasts treated with or without TGF-β1 show reduced intracellular DCN abundance in TGFβ1-induced myofibroblasts.

**Fig. S2.**
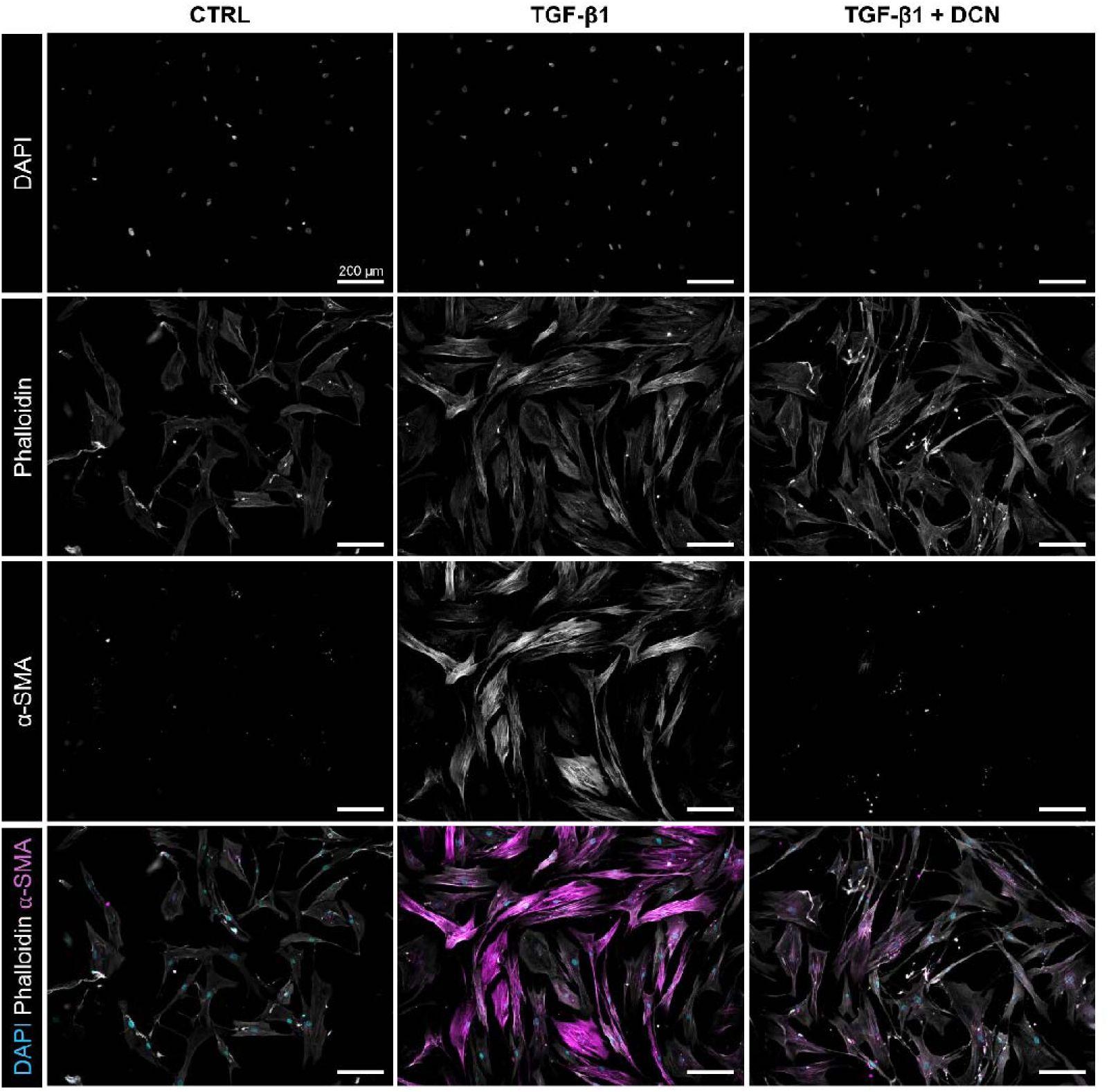
Decorin attenuates TGFβ1-induced myofibroblast activation of human VF fibroblasts. Immunocytochemistry of the myofibroblast marker α-SMA shows reduced myofibroblast activation in vitro after DCN treatment, highlighting the antifibrotic potential of restored DCN presentation.

**Fig. S3.**
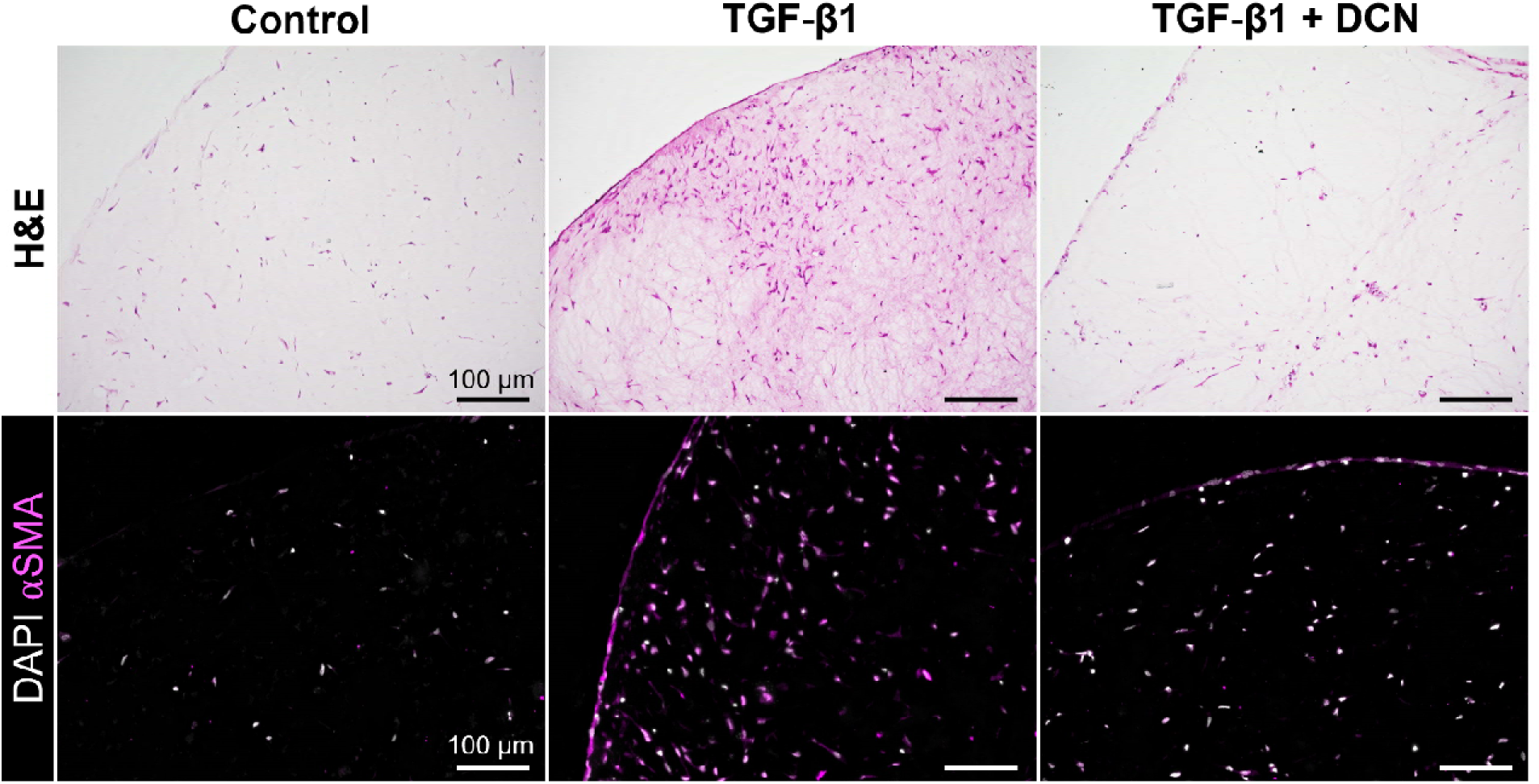
Decorin attenuates TGFβ1-induced myofibroblast activation of human VF fibroblasts in a 3D collagen hydrogel construct. Immunofluorescence of the myofibroblast marker α-SMA shows reduced myofibroblast activation in a more biomimetic 3D collagen hydrogel platform.

**Fig. S4.**
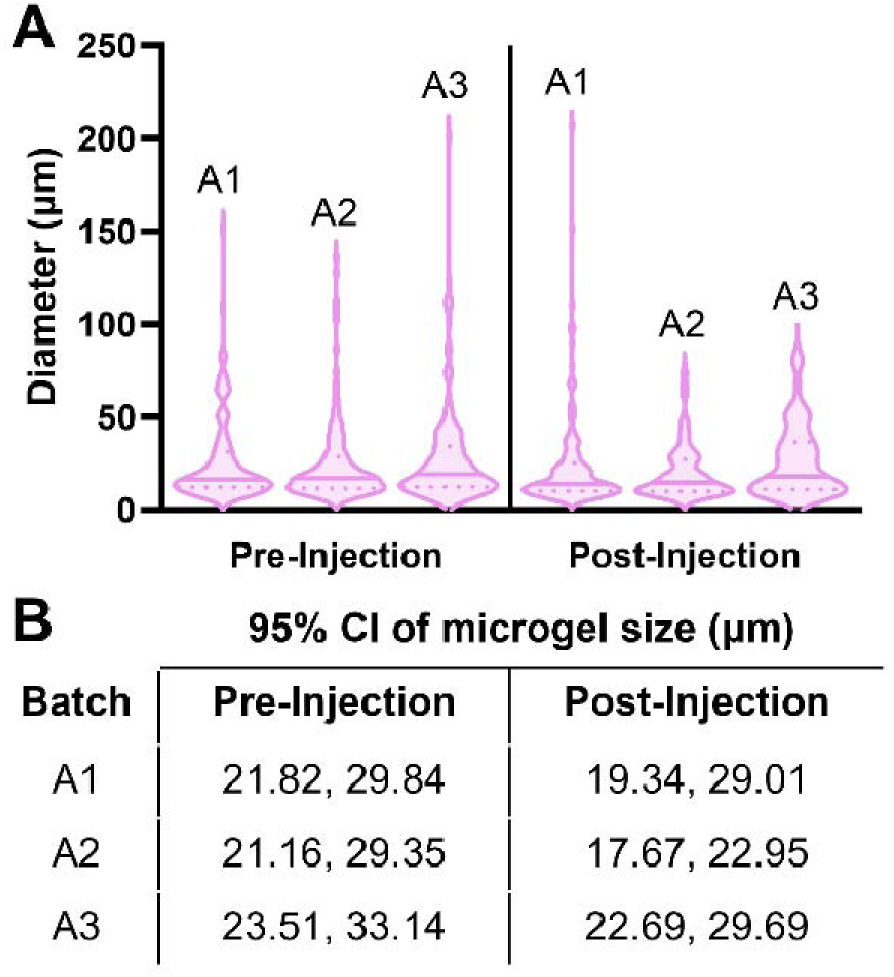
Hyaluronic acid microgels show low batch-to-batch variability. (A) Violin plot of the size distribution of hyaluronic acid microgels suggest low batch variability and negligible change in microgel size after injection with a 26G needle. (B) 95% confidence intervals (CI) of microgels from distinct batches further supports reproducibility of microgel fabrication methodology.

**Fig. S5.**
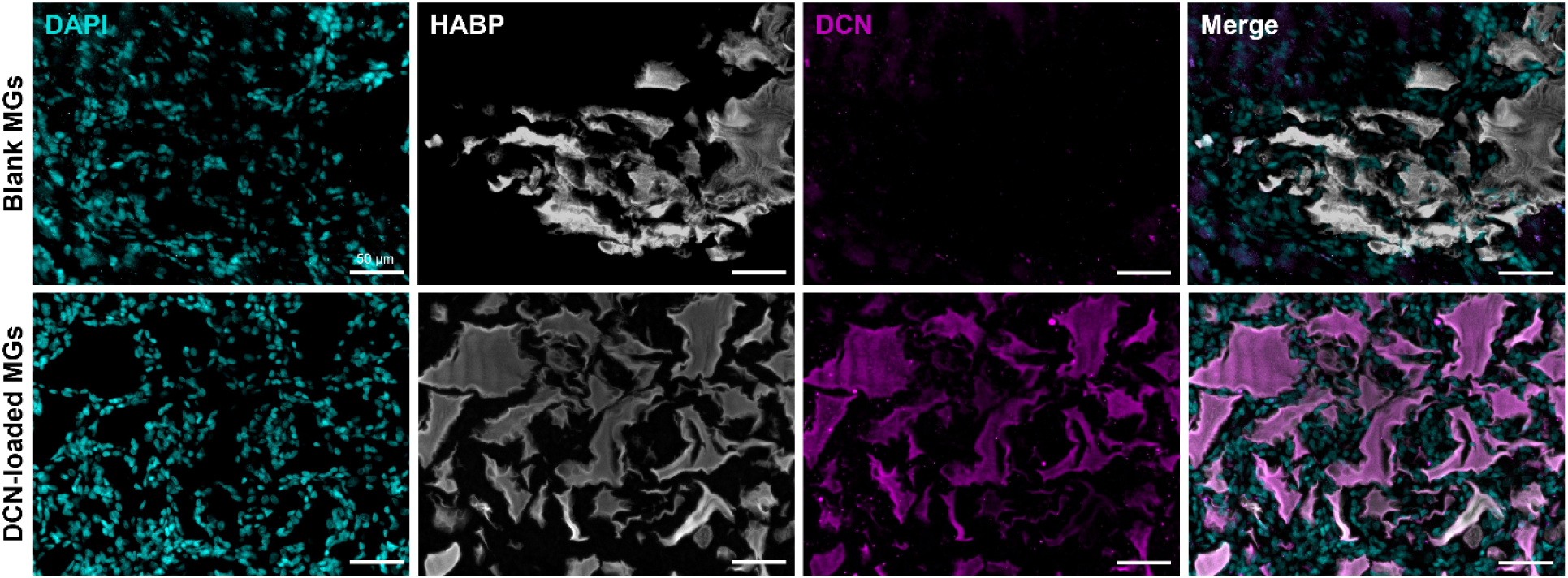
Blank and DNC-loaded microgels injected into the rat vocal fold 7 days post-injection. Microgels are easily visualized by a hyaluronan binding protein (HABP) stain in the vocal fold. DCN-loaded microgels still contain exogenous DCN protein after 7 days post-injection.

**Fig. S6.**
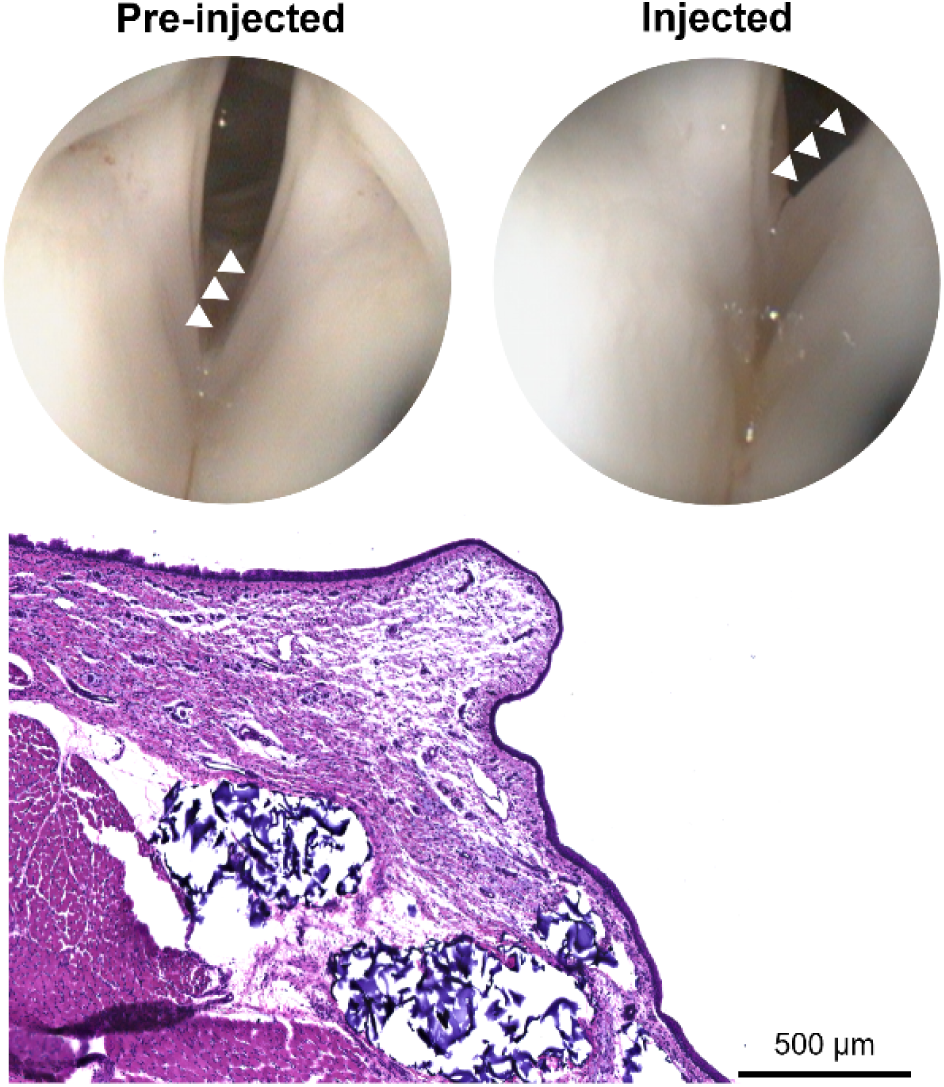
Intracordal injection of hyaluronic acid microgels in a pig model. Engineered hyaluronic acid microgels are easily injected into the vocal fold lamina propria in a porcine model to medialize the VF as shown by endoscopy and hematoxylin & eosin histology.

**Table S1.** Core matrisome differentially expressed genes in vocal fold myofibroblasts relative to unstimulated vocal fold fibroblasts.

| Annotated Gene | Annotated Matrisome Category | logFC | AveExpr | adj.P.Val |
| --- | --- | --- | --- | --- |
| COL5A1 | Collagens | 2.868714 | 11.0186034 | 1.12E-11 |
| PXDN | ECM Glycoproteins | 2.331615 | 10.28728622 | 1.11E-10 |
| SPARC | ECM Glycoproteins | 2.430636 | 12.27813059 | 1.17E-10 |
| COL1A1 | Collagens | 2.438748 | 14.77792228 | 1.57E-10 |
| DCN | Proteoglycans | -2.29279 | 10.17227255 | 2.08E-10 |
| COL4A2 | Collagens | 3.463585 | 9.91610577 | 3.57E-10 |
| DSP | ECM Glycoproteins | 2.716574 | 8.657971313 | 3.57E-10 |
| COL10A1 | Collagens | 5.159961 | 3.867611445 | 5.43E-10 |
| COL4A1 | Collagens | 4.602096 | 9.767282648 | 5.68E-10 |
| LTBP2 | ECM Glycoproteins | 3.479175 | 11.07500545 | 9.79E-10 |
| IGFBP3 | ECM Glycoproteins | 6.224382 | 10.895023 | 7.68E-09 |
| COL4A4 | Collagens | 6.075819 | 1.266850523 | 1.22E-08 |
| COMP | ECM Glycoproteins | 9.478596 | 3.340026687 | 1.70E-08 |
| ESM1 | Proteoglycans | 4.471662 | 4.475814245 | 2.29E-08 |
| FBN1 | ECM Glycoproteins | 1.595688 | 11.70251297 | 2.36E-08 |
| FN1 | ECM Glycoproteins | 2.655871 | 15.16325303 | 2.73E-08 |
| BGN | Proteoglycans | 2.66629 | 9.957809082 | 4.34E-08 |
| MATN3 | ECM Glycoproteins | 2.639081 | 2.270309007 | 4.76E-08 |
| CTGF | ECM Glycoproteins | 2.807821 | 10.69968315 | 8.83E-08 |
| COL4A3 | Collagens | 4.619464 | -0.82797163 | 1.02E-07 |
| EDIL3 | ECM Glycoproteins | 2.113924 | 7.962078354 | 1.11E-07 |
| COL7A1 | Collagens | 2.430157 | 9.781804122 | 1.62E-07 |
| HAPLN1 | Proteoglycans | 5.37355 | 7.248018298 | 1.64E-07 |
| ELN | ECM Glycoproteins | 6.079888 | 8.852841929 | 2.13E-07 |
| SPP1 | ECM Glycoproteins | 3.38486 | 0.720853851 | 2.77E-07 |
| IGFBP4 | ECM Glycoproteins | -1.56623 | 9.459792732 | 3.11E-07 |
| MATN2 | ECM Glycoproteins | -3.27935 | 3.337016864 | 3.16E-07 |
| POSTN | ECM Glycoproteins | 3.689959 | 9.566516936 | 4.09E-07 |
| COL5A2 | Collagens | 1.253039 | 10.06838858 | 4.09E-07 |
| NTN1 | ECM Glycoproteins | -4.91409 | 1.720762899 | 5.21E-07 |
| VCAN | Proteoglycans | 2.844046 | 9.607900575 | 8.97E-07 |
| SVEP1 | ECM Glycoproteins | -1.0003 | 7.840248617 | 1.58E-06 |
| FNDC1 | ECM Glycoproteins | 3.162882 | 4.32456907 | 1.78E-06 |
| IGFBP6 | ECM Glycoproteins | -1.74192 | 5.685765789 | 1.96E-06 |
| CYR61 | ECM Glycoproteins | 1.200095 | 9.304486141 | 2.18E-06 |
| SPOCK1 | Proteoglycans | 1.847653 | 8.946586748 | 2.47E-06 |
| LAMA4 | ECM Glycoproteins | -1.88509 | 7.373462429 | 2.65E-06 |
| LAMA3 | ECM Glycoproteins | -2.27689 | 2.267310647 | 2.82E-06 |
| GLDN | ECM Glycoproteins | -3.11355 | 2.541233802 | 2.99E-06 |
| SRPX | ECM Glycoproteins | -1.20003 | 6.158122825 | 3.19E-06 |
| TGFBI | ECM Glycoproteins | 3.625475 | 11.4024255 | 3.74E-06 |
| MFAP5 | ECM Glycoproteins | 3.618003 | 6.442091309 | 4.51E-06 |
| VWA5A | ECM Glycoproteins | -2.39795 | 4.388564054 | 5.85E-06 |
| THBS1 | ECM Glycoproteins | 4.064927 | 12.00533312 | 6.80E-06 |
| PODNL1 | Proteoglycans | 1.927837 | 4.269042004 | 8.40E-06 |
| FBLN5 | ECM Glycoproteins | 1.537267 | 8.922621076 | 9.71E-06 |
| COL8A1 | Collagens | 2.870402 | 7.593493263 | 1.32E-05 |
| COL8A2 | Collagens | 2.514478 | 4.85143051 | 1.49E-05 |
| IGFBP7 | ECM Glycoproteins | 1.606476 | 9.76354756 | 2.25E-05 |
| FBLN1 | ECM Glycoproteins | -1.8225 | 8.114379404 | 4.60E-05 |
| TNXB | ECM Glycoproteins | -4.75998 | 2.562110377 | 4.91E-05 |
| LAMB1 | ECM Glycoproteins | -1.16724 | 8.101257482 | 5.94E-05 |
| ZPLD1 | ECM Glycoproteins | 3.760921 | -1.03975237 | 6.39E-05 |
| MFAP3 | ECM Glycoproteins | 0.668619 | 4.686346993 | 7.06E-05 |
| HAPLN3 | Proteoglycans | 3.181272 | 3.660580023 | 0.000102 |
| COL6A6 | Collagens | -2.2731 | 2.317881968 | 0.000116 |
| COL27A1 | Collagens | -1.10355 | 4.550892833 | 0.000137 |
| FBLN7 | ECM Glycoproteins | -2.64751 | 0.279164394 | 0.000143 |
| MFAP2 | ECM Glycoproteins | 1.252572 | 6.4435062 | 0.000151 |
| IGSF10 | ECM Glycoproteins | -4.56292 | 0.275496445 | 0.000155 |
| HMCN1 | ECM Glycoproteins | 1.221292 | 7.608622097 | 0.000178 |
| VWCE | ECM Glycoproteins | -1.86898 | 2.695160973 | 0.000202 |
| PCOLCE2 | ECM Glycoproteins | -1.29584 | 4.76995832 | 0.000231 |
| NOV | ECM Glycoproteins | -2.38166 | 4.350879405 | 0.000259 |
| LAMB3 | ECM Glycoproteins | -1.09828 | 3.850909351 | 0.000276 |
| COL18A1 | Collagens | -1.07269 | 6.349259909 | 0.000296 |
| COL16A1 | Collagens | 0.688077 | 8.778067322 | 0.000346 |
| COL1A2 | Collagens | 0.853659 | 13.6671557 | 0.000374 |
| COL9A2 | Collagens | -2.18384 | 1.566718291 | 0.000461 |
| COL15A1 | Collagens | 5.01604 | 5.118189976 | 0.000524 |
| PRELP | Proteoglycans | -2.49969 | 2.452090718 | 0.000607 |
| CTHRC1 | ECM Glycoproteins | 1.495283 | 8.680351543 | 0.000712 |
| SMAP2 | ECM Glycoproteins | -0.754 | 4.98018292 | 0.000717 |
| LAMC1 | ECM Glycoproteins | 0.631335 | 10.77109633 | 0.000995 |
| COL12A1 | Collagens | 0.857736 | 9.74493337 | 0.001003 |
| SMOC1 | ECM Glycoproteins | 0.949571 | 2.91500979 | 0.001242 |
| TNFAIP6 | ECM Glycoproteins | 1.470475 | 4.516265749 | 0.001506 |
| COL3A1 | Collagens | 1.093301 | 11.19557159 | 0.001815 |
| PXN | ECM Glycoproteins | 0.368164 | 8.312110815 | 0.002261 |
| POM121 | ECM Glycoproteins | 0.296225 | 5.379070562 | 0.004729 |
| ASPN | Proteoglycans | 1.782556 | 5.941091151 | 0.004913 |
| LAMA2 | ECM Glycoproteins | -0.82339 | 6.246620889 | 0.005049 |
| THBS2 | ECM Glycoproteins | 1.29066 | 9.417562178 | 0.00599 |
| MBP | Proteoglycans | -1.89225 | 2.76668503 | 0.006087 |
| LAMC2 | ECM Glycoproteins | 0.900729 | 2.838145685 | 0.007779 |
| LUM | Proteoglycans | -0.5888 | 7.620417042 | 0.008091 |
| VWDE | ECM Glycoproteins | -1.35338 | -1.06525257 | 0.00846 |
| EMILIN2 | ECM Glycoproteins | -1.48768 | 2.985052862 | 0.009725 |
| LTBP4 | ECM Glycoproteins | -1.68347 | 6.229997854 | 0.009859 |
| NID2 | ECM Glycoproteins | -1.07086 | 6.42281128 | 0.01086 |
| LAMA5 | ECM Glycoproteins | -1.74988 | 4.942330005 | 0.01095 |
| FBLN2 | ECM Glycoproteins | -1.35094 | 7.691113367 | 0.01165 |
| AGRN | ECM Glycoproteins | 0.721691 | 7.710235484 | 0.011784 |
| BMPER | ECM Glycoproteins | -1.43166 | 2.473001504 | 0.012186 |
| CRIM1 | ECM Glycoproteins | 0.824151 | 8.913075659 | 0.012578 |
| HSPG2 | Proteoglycans | 0.830649 | 9.977843432 | 0.014669 |
| COL19A1 | Collagens | -1.54848 | -0.429575896 | 0.015637 |
| VWF | ECM Glycoproteins | -1.67906 | 1.0217903 | 0.018025 |
| COL14A1 | Collagens | -3.25232 | 3.626318865 | 0.020451 |
| LTBP3 | ECM Glycoproteins | -0.87359 | 6.518265396 | 0.02106 |
| SLIT2 | ECM Glycoproteins | -0.67978 | 5.372168499 | 0.02142 |
| PODN | Proteoglycans | -0.96549 | 6.406055656 | 0.021627 |
| OGN | Proteoglycans | -3.65682 | 0.184102093 | 0.024556 |
| FGL2 | ECM Glycoproteins | -3.995 | -0.953349513 | 0.026185 |
| COL5A3 | Collagens | 1.132404 | 3.987406425 | 0.035874 |
| GAS6 | ECM Glycoproteins | 0.641718 | 10.52968547 | 0.047606 |
| AEBP1 | ECM Glycoproteins | 0.731755 | 9.555305253 | 0.047999 |
| VWA9 | ECM Glycoproteins | 0.220892 | 4.58124939 | 0.052747 |
| SRGN | Proteoglycans | 1.3089 | 4.540498406 | 0.05468 |
| THSD4 | ECM Glycoproteins | -0.45877 | 5.964724247 | 0.054995 |
| NRP2 | ECM Glycoproteins | 0.7371 | 5.552078392 | 0.058191 |
| LTBP1 | ECM Glycoproteins | 0.495561 | 9.376674005 | 0.063884 |
| COL11A1 | Collagens | 5.008814 | 2.733802216 | 0.064059 |
| TNC | ECM Glycoproteins | 1.465523 | 7.568830981 | 0.069472 |
| SRPX2 | ECM Glycoproteins | 0.675526 | 6.06577457 | 0.07238 |
| ECM1 | ECM Glycoproteins | -0.56886 | 7.772805894 | 0.077922 |
| COL6A1 | Collagens | -0.58832 | 11.17696626 | 0.082751 |
| SPON1 | ECM Glycoproteins | -2.00433 | -1.843947214 | 0.085563 |
| NTNG2 | ECM Glycoproteins | -0.58742 | 2.428708377 | 0.104539 |
| COL4A6 | Collagens | -2.01423 | -1.675095249 | 0.106904 |
| COL13A1 | Collagens | -0.95555 | 3.573629273 | 0.110823 |
| MXRA5 | ECM Glycoproteins | -1.69559 | 4.599851433 | 0.111284 |
| MFGE8 | ECM Glycoproteins | 0.6572 | 8.806075949 | 0.113614 |

**Table S2.**
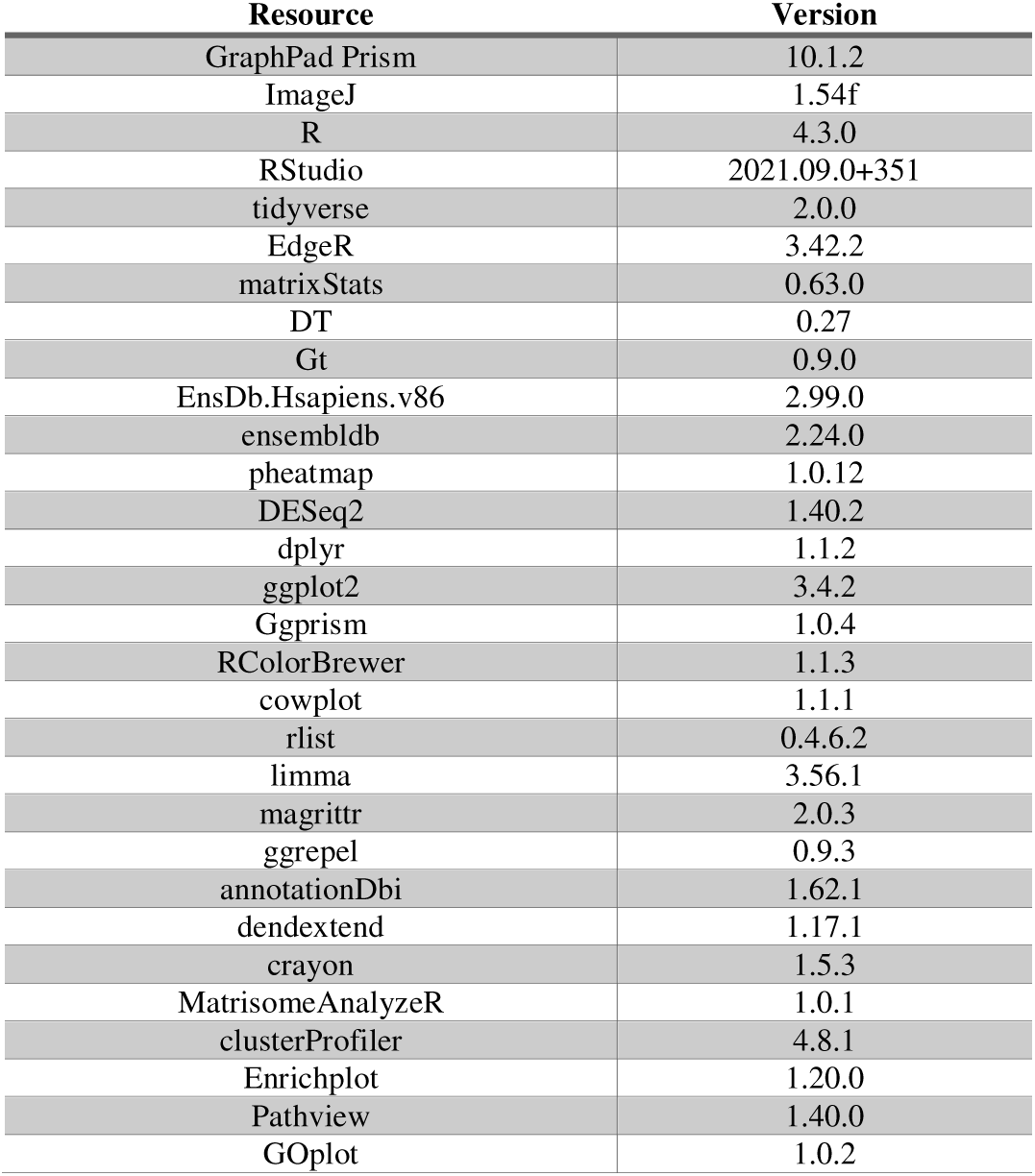
Resources, software, and R packages used for image analysis or bulk transcriptomics.

**Table S3.**
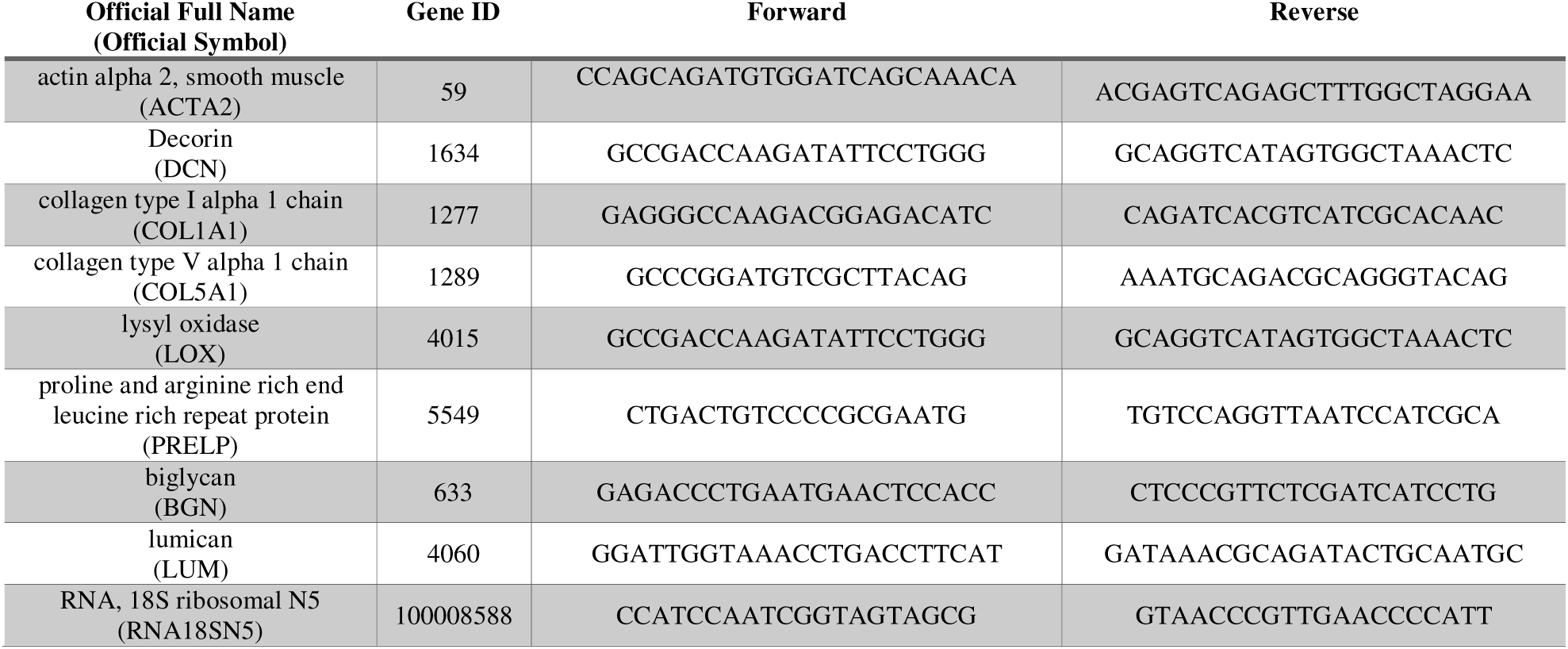
List of oligonucleotide primers.

**Table S4.** List of antibodies.

| Antibody/Binding Protein | Dilution | Supplier (Cat No.) |
| --- | --- | --- |
| Rabbit IgG anti-decorin,<br>polyclonal antibody | 1:200 | Proteintech (14667-1-AP) |
| Mouse IgG2a,kappa anti-α-<br>smooth muscle actin<br>monoclonal antibody (1A4) | 1:500 | Invitrogen (14-9760-82) |
| Hyaluronan Binding Protein<br>(HABP) (Biotin conjugate) | 1:60 | AMSBio LLC (AMS.HKD-BC41) |
| Goat anti-rabbit IgG (H+L),<br>F(ab')<sub>2</sub> fragment (Alexa Fluor®<br>488 conjugate) | 1:750 | Cell Signaling Technology (4412S) |
| Streptavidin, Alexa Fluor® 488 | 1:500 | ThermoFisher Scientific (S32354) |

| conjugate |  |  |
| --- | --- | --- |
| Goat anti-mouse IgG (H+L),<br>F(ab')<sub>2</sub> fragment (Alexa Fluor®<br>647 conjugate) | 1:750 | Cell Signaling Technology (4410S) |
| Goat anti-rabbit IgG (H+L),<br>F(ab')<sub>2</sub> fragment (Alexa<br>Fluor® 647 conjugate) | 1:750 | Cell Signaling Technology (4414S) |

